# Experts, but not novices, exhibit StartReact indicating experts use the reticulospinal system more than novices

**DOI:** 10.1101/841791

**Authors:** Brandon M. Bartels, Maria Jose Quezada, Vengateswaran J. Ravichandran, Claire F. Honeycutt

## Abstract

Motor skill acquisition utilizes a wide array of neural structures; however, few articles evaluate how the relative contributions of these structures shift over the course of learning. Recent evidence from rodents and songbirds suggests there is a transfer from cortical to subcortical structures following intense, repetitive training. Evidence from humans indicate that the reticulospinal system is modulated over the course of skill acquisition and may be a subcortical facilitator of learning. The objective of this study was to evaluate how reticulospinal contributions are modulated by task expertise. Reticulospinal contributions were assessed using StartReact (SR). We hypothesized that expert typists would show SR during an individuated, keystroke task but SR would be absent in novices. Expert (75.2 ± 9.8 WPM) and novice typists (41.6 ± 8.2 WPM) were evaluated during an individuated, keystroke movements. In experts, SR was present in all fingers (except the middle) but was absent in novices (except the little). Together, these results suggest that experts use reticulospinal contributions more for movement than novices indicating that the reticular formation becomes increasingly important for movement execution in highly trained, skilled tasks even those that require individuated movement of the fingers.

## Introduction

Motor skill acquisition, the process by which movements are honed and refined to become faster and more accurate, utilizes a wide array of neural structures – cortical, subcortical, cerebellar, and brainstem – which interact during skill acquisition. While there has been significant evaluation of the role of these neural structures in motor learning, few articles evaluate how the relative contributions of these structures shift over the course of skill acquisition. A recent study demonstrated that there is a transfer from the use of cortical to subcortical structures following intense, repetitive training (Kawai et al. 2015). Kawai et al. demonstrated that the initial learning of a task requires substantial input from the cortex; however following training, rats with bilateral motor cortical lesions were still able to perform a sequence of precisely timed lever presses, i.e. if sufficiently trained prior to lesion, rats can perform lever presses in the absence of the motor cortex indicating a shift to more heavy involvement of subcortical structures. This phenomenon is not isolated to rats. Similarly, non-human primates show dexterous movement following large, gray-matter cortical lesion (Darling et al. 2018) and songbirds with bilateral forebrain cortical lesions can also perform songs with appropriate frequency and amplitude modulations (Andalman and Fee 2009; Bottjer et al. 2006; Turner and Desmurget 2010). Interestingly, songbirds can only perform songs based on their developmental or expertise level when the lesioning occurred. For example, young, novice birds can only perform rudimentary songs but older, expert songbirds can perform more complex songs of larger variation. Importantly, neither rats nor songbirds can learn new tasks or songs if lesioning occurs prior to learning. This highlights a critical role for both the cortical and subcortical systems during learning but also notes that following learning subcortical systems become increasingly important for execution of learned tasks.

While several subcortical structures may be involved in the transfer from cortical to subcortical structures (cerebellum, basal ganglia, red nucleus, superior colliculus), a handful of studies indicate that the reticular formation may be an important facilitator. Evidence from humans shows that reticulospinal contributions are modulated during intense, repetitive training. Direct measurement of the reticulospinal system in humans is not feasible; thus the StartReact (SR) response is utilized to assess reticulospinal contributions. The use of SR to investigate the reticular formation in humans is predicated on animal work (Davis et al. 1982; Davis and Gendelman 1977; Groves et al. 1974; Hammond 1973); however, strong evidence exists that the role of the reticular formation during SR is maintained in humans. Patients with heredity spasticity disorder (HSP) have selective corticospinal tract damage but intact reticulospinal pathways. In these patients, SR is intact and further shows no evidence of deficit or delay (Nonnekes et al. 2014b). Additionally, SR remains intact following cortical lesion in stroke survivors (Honeycutt et al. 2014; Honeycutt and Perreault 2012a; Marinovic et al. 2016) indicating that SR does not rely on the cortex or corticospinal projection for execution in humans.

Prior studies show that the SR response is modulated during intense, repetitive training indicating that the reticulospinal system may be involved in skill acquisition. When the SR response is monitored over the course of a 10-day training regimen of an index finger abduction, SR absent on Day 1 but develops by Day 10 indicating increasing reticulospinal contributions to movement execution (Kirkpatrick et al. 2018). In addition, the SR response shows differential response characteristics before and after training (Maslovat et al. 2009, 2011). Specifically, SR is responsive to training decreasing within-subject kinematic variability. Maslovat et. al.’s 2011 study was the first to show that learning present during voluntary movements is represented during corresponding SR movements, i.e. as voluntary movements become less variable, SR movements also become less variable.

Together, these studies indicate that the reticulospinal system is modulated over the course of skill acquisition and may be an important facilitator of learning. Still, previous studies evaluated short-term effects with no consideration for long-term retention such as the task expertise similar to that of advanced, expert songbirds. The objective of this study was to evaluate if reticulospinal contributions are modulated with task expertise (long-term skill retention). We evaluate typing expertise because 1) expertise is easily quantified with typing speeds, 2) typing has been used extensively to evaluate motor learning and skill acquisition (Cannonieri et al. 2007; Chapman 1919; Hill 1934; Hill et al. 1913; Sternberg et al. 1978), and 3) typing represents a common task of significance to many adults. As it can take months and arguable years to become an expert typist, we evaluated 2 groups of subjects categorized as expert or novice typists based on a three-minute typing test. We hypothesized that experts would show SR during an individuated, keystroke task demonstrated by a difference between Startle+ (presence of startle) and Startle-(absence of startle) trials but SR would be absent in novices. If true, it would indicate that the reticulospinal system is utilized more by experts than novices suggesting an important role of the reticulospinal system during skill acquisition and long-term skill retention.

## Methods

This study was approved by Institutional Review Board STUDY00002440 under Arizona State University. Subjects were informed of all potential risks prior to participation in the study and verbal/written consent was obtained.

### Subjects

Seventeen neurologically unimpaired, right-handed individuals (9 Male, 8 Female; Age: 20.7 ± 0.9 years) were used for this study. Subjects were categorized as expert or novice typists based on the result of a three-minute typing test. The average number of correct words typed per minute was used to account for speed and accuracy. An unpaired t-test confirmed that the expert population (75.2 ± 9.8 WPM) scored higher on the test than the novice population (41.6 ± 8.2 WPM) (t_stat_ = 7.44, P < 0.001).

### Data acquisition

EMG data were collected at 3000 Hz with Ag/AgCl bipolar surface electrodes [MVAP Medical Supplies, Newbury Park, CA], two Bortec AMT-8 amplifiers [Bortec Biomedical Ltd., Canada], and a 16-bit data acquisition system (NI USB-6363, National Instrumentation, Austin, TX). The amplifiers (gain = 3000) had an internal bandpass filter set at 10-1000 Hz. To record finger extension and flexion, EMG was collected from the Abductor Pollicis Brevis (APB - thumb), Extensor Digitorium Communis (ED2-index, ED3-middle, ED4-ring), Extensor Digiti Minimi (EDM - little), and Flexor Digitorum Superficialis (F2/3-index/middle, F4-ring, F5-little) muscles using a protocol established in the literature for evaluating individuated finger movements (Leijnse et al. 2008). To monitor startle, the right and left Sternocleidomastoid (RSCM, LSCM) muscles were recorded (Carlsen et al. 2011; Leow et al. 2018). In addition to EMG, the keystroke was monitored using the change in voltage from the instrumented keyboard.

At the beginning of each trial, the subject positioned their right hand over the J (index finger), K (middle finger), L (forth finger), and; (little finger) keys because expert typists are taught to use these keys as the “home row.” Subjects were instructed to perform a single keystroke, of the specified key (e.g. J), following a series of auditory tones. The first tone, a soft acoustic stimulus of 80-dB, informed the subject to start planning the task (‘GET READY’). Between 2.5-3.5 seconds later, the subject was provided with a ‘GO’ cue of a soft, 80-dB sound. During 33% of trials the ‘GO’ was randomly replaced with a loud, 115-dB acoustic stimulus which is known to induce a SR response (Carlsen et al. 2004; Valls-Solé et al. 1995, 1999). Each subject completed a total of 225 trials (45 trials for each key/finger). The order of keys was randomized into blocks of 15 key strokes during which the subject pressed a single key (e.g. J). Subjects were given the opportunity to rest after each set of 15.

### Data processing

EMG data were rectified and smoothed in Matlab (R2017b) using a 10-point moving average. Muscle onset latency was evaluated using a custom Matlab script that selected the first instance the EMG signal achieved greater than three times the standard deviation of background activity. Background activity was defined as the average of a 500 ms period prior to the GO cue. This selected onset was then visually inspected by a researcher blinded to all independent variables. Keystroke onset latency was also detected automatically with a computer and visually verified by the experimenter blinded to the trial type. LSCM and RSCM muscles were used to separate loud, acoustic trials into Startle+ (presence of startle) and Startle-(absence of startle). Startle+ was defined as presence of an SCM response prior to 120ms in either the LSCM or RSCM. The SR effect was evaluated by comparing Startle+ to Startle-trials during loud, acoustic stimulus trials to account for intensity related effects (Carlsen et al. 2007, 2009; Honeycutt et al. 2013; Leow et al. 2018). If the onset latencies were different, SR was present. As our experimental design required the presence of both Startle+ and Startle-trials, we used a loud, acoustic stimulus of 115dB which is lower than most SR publications which report Startle during most trials where a loud, acoustic stimulus is present. Conditions where at least one Startle+ or Startle-was not present (6.2%) were not included. Trials in which the user failed to press the key, pressed multiple keys, or pressed the key too late (i.e. EMG latency > 300ms, keystroke latency > 350ms) were eliminated from analysis (4.01% of trials). Additionally, one subject was eliminated from analysis as 63% of their trials had keystroke onsets later than 350ms.

### Statistical analysis

Startle+ and Startle-EMG and keystroke onset latencies were compared using R statistical software (v3.4.2). Startle+ and Startle-EMG and keystroke onset latencies for both populations were compared using a 3-way, generalized linear mixed effects model (GLMM) that did not assume equal variance (Cohen 1988). Condition (Startle+ or Startle-) and Expertise (expert or novice) and Muscle (e.g. APB) were the independent variables, onset latencies (EMG and keystroke latencies) were the dependent variables, and subjects were treated as a random factor. A confidence interval of 95% was used to classify statistical significance.

As the hypothesis was that novices would have no difference between Startle+ and Startle-, EMG and keystroke onset latencies, novices were further compared with a two one-sided T-tests (TOST) to test the null hypothesis that the Startle+ and Startle-latencies were different. The TOSTER package in R was executed. As is convention for TOST, we used a 90% confidence interval (5% for each side). We conservatively used the smaller standard deviation of the two distributions as the equivalence bound. First, to prevent an increase in Type I error, that occurs when running this analysis independently for each muscle, latencies were normalized to their corresponding maximum latency for each muscle/condition prior to conducting a test of equivalence. This is reported as the group results. However, this can increased Type II errors, thus to minimize Type II errors and given the repeated measures design, we also used the 80th percentile from a one-sided Chi-squared distribution (γ = 0.2) as the estimate of measurement precision (Limentani et al. 2005). Matching this, we used the 80th percentile of distribution (1.3 * smallest standard deviation) as the equivalence bounds for testing equivalence of the distributions. Both group and individual muscle results are reported. Shapiro-Wilks test on the normalized latencies confirms the normality of both the Startle-(P ~= 0) and Startle+ (P ~=0) conditions.

To test for differences in the probability of startle (Count of Startle+ Trials / Count of Startle+ and Startle-Trials), a GLMM with population (expert and novice) and finger as independent variables was executed. All data in the results section is presented with marginalized means and standard errors. Finally, we used a Brown-Forsythe test to test how variability differed between experts and novices.

## Results

The 3-way GLMM found the effect of Condition (F_2,5801_ = 37.1 P < 0.001) and Muscle (F_1,5800_ = 7.29; P = 0.007) but not Expertise (F_1,20_ = 1.16; P = 0.29). However, there was a significant interaction effect for Condition:Expertise (F_2,5801_ = 4.05; P = 0.017) and Muscle:Expertise (F_1,5800_ = 6.24; P = 0.012) that was further explored with post-hoc testing. Startle+ and Startle-EMG and keystroke onset latencies were different in experts showing an SR effect (Fig. 1). In experts, the onset latency of Startle+ trials was faster than Startle-trials in the thumb (APB: Δ = 9.11, P = 0.041, *Keystroke*: Δ = 13.98, P = 0.0004), index finger (*ED2*: Δ =14.01, P = 0.016; *F2*: Δ = 10.46, P = 0.013; *Keystroke*: Δ =11.86, P = 0.004), ring finger (*ED4*: Δ = 30.66, P = 0.005; *F4*: Δ = 17.62, P = 0.0047; *Keystroke*: Δ =19.61, P = 0.011), and the little finger (*EDM*: Δ = 21.02, P = 0.002; *F5*: Δ = 18.06, P = 0.0034; *Keystroke*: Δ = 16.39, P = 0.0007). However, Startle+ and Startle-onset latencies were not different in the middle finger (*ED3*: Δ = −2.34, P = 0.97; *F3*: Δ = 1.55, P = 0.79 *Keystroke*: Δ = 0.79, P = 0.63).

**Figure 1.**
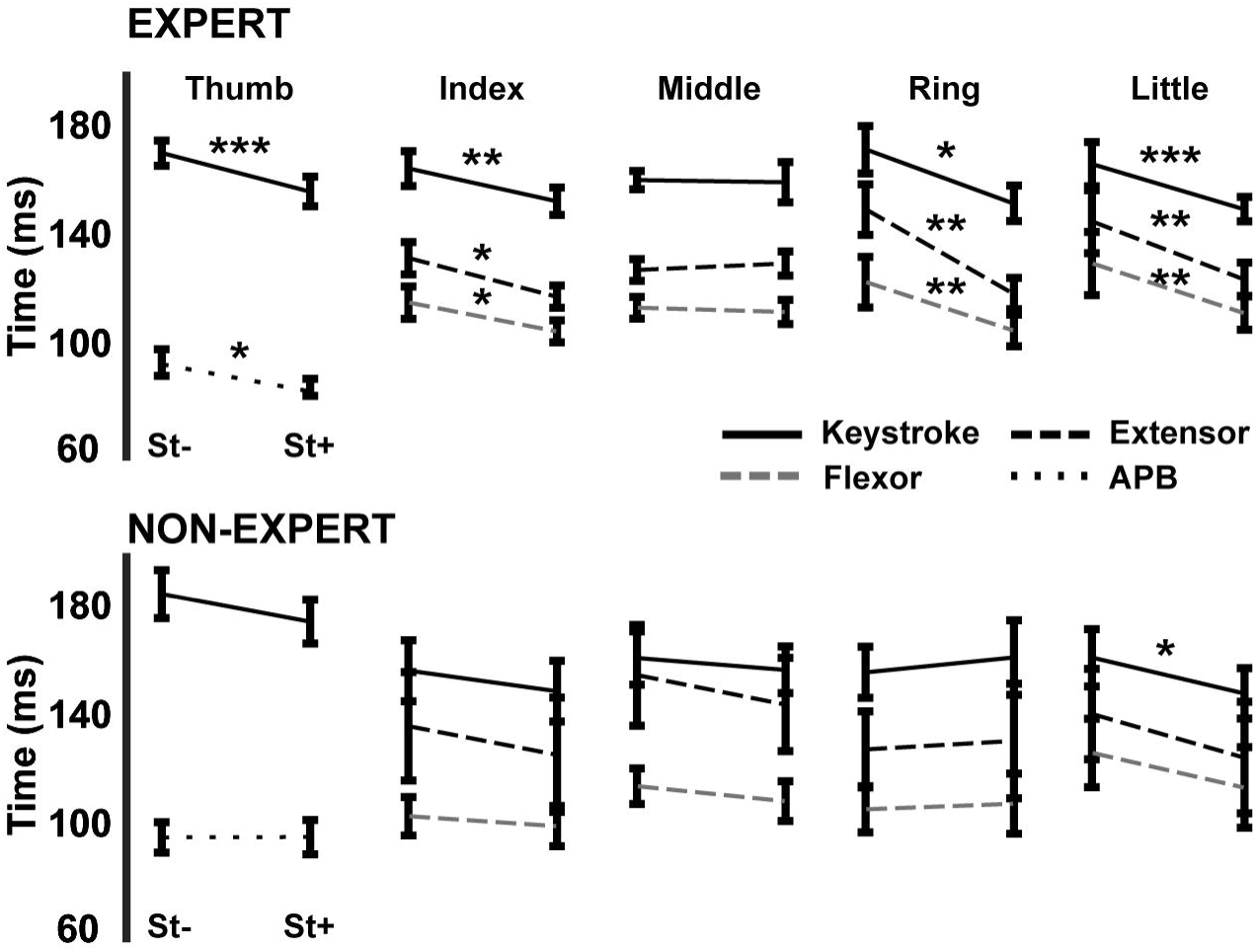
Startle+ and Startle-trials are compared between experts and novices. Onset Latency of the keystroke (solid black) as well as extensor (dashed black) and flexor (dashed gray) muscles in experts (top) and novices (bottom) are displayed for Startle+(right) and Startle-(left) trials. Stars represent a difference between Startle+ and Startle-(* = P < 0.05, ** = P < 0.01, and *** P< 0.001).

Onset latencies between Startle+ and Startle-trials were not different in novices except in the little finger (and not robustly) indicating no SR effect. Onset latencies of Startle+ trials were not different from Startle-trials in the thumb (*APB*: Δ = −0.13, P = 0.42; *Keystroke*: Δ = 9.99, P = 0.094), index (*ED2*: Δ = 10.30, P = 0.295; *F2*: Δ = 3.58, P = 0.204; *Keystroke*: Δ = 7.40, P = 0.30), middle (*ED3*: Δ = 10.61, P = 0.26; *F3*: Δ = 5.39, P = 0.24; *Keystroke*: Δ = 4.28, P = 0.43), and ring (*ED4*: Δ = −3.06, P = 0.46; *F4*: Δ = −2.00, P = 0.32; *Keystroke*: Δ = −5.37, P = 0.10). Startle+ and Startle-onset latencies were different for the little finger in the keystroke (Δ = 12.94, P = 0.03), but did not reach significance in the muscle responses (EDM: Δ = 15.75, P = 0.07; *F5*: Δ = 12.55, P = 0.23).

TOST further confirmed that the difference between Startle+ and Startle-in novices did not reach significance. When all values were considered the null hypothesis that Startle+ and Startle-trials are different was rejected (t_775.281_ = −11.640, P = <0.0001). When muscles were compared individually, the null hypotheses were also rejected (all t < −1.3 and all P = < .005). The overall difference between the novice Startle+ and Startle-normalized onset latencies was 9.4% (90% confidence interval: 7.3, 11.4).

Differences associated with startle between experts and novices were not the result of differences in the probability of startle. The probability of evoking startle was not statistically different between populations (F_1,75_ = 0.01, P = 0.92) or between fingers (F_4,75_ = 0.44, P = 0.78). Additionally, there was no difference in interaction between fingers and population (F_4,75_ = 0.13, P = 0.97). The probability of startle for the expert and novice populations, when averaged between fingers, was 34.97 ± 4.93% and 36.62 ± 4.65% respectively. The group variability between experts and novices did not reach significance (F_1,1717_ = 3.011; p = 0.08). However, individual muscle comparisons showed that the variability was higher in the APB (F_1,178_ = 9.16; P = 0.002) and ED3 (F_1,197_; P < 0.001) muscles.

## Discussion

### Summary

The objective of this study was to evaluate if reticulospinal contributions are modulated with task expertise (long-term skill retention). As the presence of an SR response indicates the presence of reticulospinal contributions (Baker and Perez 2017; Carlsen et al. 2009; Honeycutt et al. 2013), we hypothesized that experts would show SR during an individuated, keystroke task demonstrated by a difference between Startle+ (presence of startle) and Startle-(absence of startle) trials but SR would be absent in novices. We found that experts have an SR effect in all fingers (except the middle) but the SR effect was absent in novices. These results suggest that experts use reticulospinal contributions more for typing-like movements than novices indicating that the reticular formation, along with other subcortical structures, are increasingly important for movement execution in highly trained, skilled tasks even those that require individuated finger movement.

### Movement execution in experts vs. novices

Our results add to the growing list of studies demonstrating that the neural control of movement is altered over the course of skill acquisition specifically highlighting the importance of subcortical structures in the execution of highly trained tasks. The presence of SR during individuated finger movement of experts (but not novices) indicates that the reticulospinal system becomes increasingly involved in the execution of a task as it is learned, even during individuated movements of the fingers. Conversely, the absence of SR in novices suggests that the reticular formation is not used (or used less) during the execution of novel or un-practiced tasks – particularly of the fingers. It is notable that a subset of the muscles in the novice population had increased variability. While most did not reach statistical significance, the thumb (APB) and middle finger (ED3 but not F3) showed higher variability in novices compared to experts. This is consistent with the literature showing that during learning, subjects display exploratory patterns of behaviors (Fee and Goldberg 2011; Neuringer et al. 2000; You and JM 2019) that result in increased variability; however, over time neural activity consolidates to a refined pattern of behavior (Churchland et al. 2012; Kao et al. 2015; Sadtler et al. 2014). The initial high variability during learning may also be linked to an inability to adequately “prepare” a finely tuned motor program. It has been argued that SR is tied to preparation level (Marinovic and Tresilian 2016). If we define preparation as the capacity to load a specific, highly trained, feedforward command, then novices, who are using exploratory patterns that differ on a trial-by-trial basis, would not able to prepare a highly trained motor program. Thus, novices likely are using many different neural substrates during task execution, possibly even subcortical substrates, over the course of learning but the increased variability present when observing novice behavior in certain muscles makes differentiation of these substrates more difficult. Future studies should further evaluate how preparation level might differ over the course of motor learning and how SR may be used as a quantitative method to evaluate task expertise.

### Startle as a behavioral indicator of motor learning

SR has been previously suggested to be a measurable behavioral indicator of motor learning because it exhibits changes during motor learning (Kirkpatrick et al. 2018; Maslovat et al. 2008, 2009, 2011, 2012). The data here indicate that SR can also distinguish task expertise which might prove useful in rehabilitation by determining when a task has been successfully “re-learned”. SR is readily present in patient populations even enhancing functional performance in some cases (Baker and Perez 2017; Fletcher and Jon 2014; Honeycutt et al. 2013, 2015; Honeycutt and Perreault 2012b; Marinovic et al. 2016; Marinovic and Tresilian 2016; Nonnekes et al. 2014a, 2014b, 2015; Rahimi and Honeycutt 2017; Serranová et al. 2012). Thus, future studies should explore if the presence of SR during a movement relates to retention of therapy in more acute populations.

### Motor independence and the lack of SR in the middle finger

A surprising result was the absence of SR in the middle finger of experts. The absence of SR in the middle finger may be related to motor independence. Individuated finger movements have different levels of “motor independence.” There are 36 muscles used for manipulation of the thumb and fingers (5 digits), all of which act synergistically (Santello et al. 2013) in coordinated neuromuscular patterns (Schieber and Santello 2004). This synergistic activation leads to a coupling of fingers when trying to perform individuated movements (Fish and Soechting 1992; Hager-Ross and Schieber 2000). For example, the middle and ring fingers have the lowest levels of motor independence (Hager-Ross and Schieber 2000) resulting in coupling of motor actions when a subject is asked to move them independently. Lack of motor independence may explain the absence of SR in the middle finger but does not explain the presence of SR in the ring finger which also has low motor independence? Unlike the middle finger, the ring finger has significant and overlapping neural correlates with the little finger (Hager-Ross and Schieber 2000). The little finger has a robust SR response in experts and further exhibits a small response in non-experts. Thus, the little finger may “rescue” the ring finger’s SR response. Further evaluation is needed to explore this possibility as well as evaluate if SR is sensitive to motor independence.

## Acknowledgments

This work was supported by Arizona State University Fulton Undergraduate Research Initiative Arizona State, and University Grand Challenge Scholars Program.

